# Warm periods in repeated cold stresses protect *Drosophila* against ionoregulatory collapse, chilling injury, and reproductive deficits

**DOI:** 10.1101/2020.02.04.934265

**Authors:** Mahmoud I. El-Saadi, Marshall W. Ritchie, Hannah E. Davis, Heath A. MacMillan

## Abstract

In many insects, repeated cold stress, characterized by warm periods that interrupt cold periods, have been found to yield survival benefits over continuous cold stress, but at the cost of reproduction. During cold stress, chill susceptible insects like *Drosophila melanogaster* suffer from a loss of ion and water balance, and the current model of recovery from chilling posits that re-establishment of ion homeostasis begins upon return to a warm environment, but that it takes minutes to hours for an insect to fully restore homeostasis. Following this ionoregulatory model of chill coma recovery, we predicted that the longer the duration of the warm periods between cold stresses, the better a fly will recover from a subsequent chill coma event and the more likely they will be to survive, but at the cost of fewer offspring. Here, female *D. melanogaster* were treated to a long continuous cold stress (25 h at 0°C), or experienced the same total time in the cold with repeated short (15 min), or long (120 min) breaks at 23°C. We found that warm periods in general improved survival outcomes, and individuals that recovered for more time in between cold periods had significantly lower rates of injury, faster recovery from chill coma, and produced greater, rather than fewer, offspring. These improvements in chill tolerance were associated with mitigation of ionoregulatory collapse, as flies that experienced either short or long warm periods better maintained low hemolymph [K^+^]. Thus, warm periods that interrupt cold exposures improve cold tolerance and fertility in *D. melanogaster* females relative to a single sustained cold stress, potentially because this time allows for recovery of ion and water homeostasis.

## Introduction

Insects exposed to cold temperatures below their chill coma onset temperature (CCO) enter chill coma, a narcosis-like state characterized by a loss of nerve and muscle function (Andersen et al., 2015; Hazell and Bale, 2011;MacMillan and Sinclair, 2011). In many insects, including the model organism, *Drosophila melanogaster*, cold-induced disruption in ion homeostasis occurs through suppression of active ion transport across the renal epithelia and leak of K^+^ down its concentration gradient from tissues (Overgaard and MacMillan, 2017). This failure of organismal ion balance causes hemolymph volume to decrease as gut water volume increases, and concentrates K^+^ in the remaining hemolymph (MacMillan et al., 2015a). When a chill-susceptible insect remains at unfavourable low temperatures for a prolonged period, it can start to accumulate chilling injuries that manifest as motor defects or an inability to recover the ability to move or continue development upon rewarming (Koštál et al., 2006, 2004; MacMillan and Sinclair, 2011). Damage to the muscles, in particular, is thought to be caused by hemolymph hyperkalemia that causes a depolarization of muscle membrane potentials, dysregulation of Ca^2+^ balance, and activation of apoptotic signalling cascades (Andersen et al., 2017; Bayley et al., 2018; MacMillan et al., 2015b).

If a cold exposure is mild or brief enough such that an insect survives without extensive chilling injury, chill coma can be reversed, and chill coma recovery time (CCRT), like chill coma onset or a survival assessment following a cold stress, is a common measure of insect cold tolerance (Anderson et al., 2005; Gibert et al., 2001; Milton and Partridge, 2008). When an insect is rewarmed, ionoregulatory function is largely restored and hemolymph [K^+^] begins to return to near-baseline levels (Findsen et al., 2013; MacMillan et al., 2014; MacMillan et al., 2012). This recovery of ion balance upon rewarming is most likely due to an influx of K^+^ into the muscles and other tissues (MacMillan et al., 2014), as well as increased ion transport at the renal epithelia (MacMillan et al., 2012, 2015a). During this process, hemolymph [K^+^] can be restored in as little as 10-15 min in crickets and locusts, but full recovery of water balance can take far longer (e.g. 2 h) (MacMillan et al., 2014; MacMillan et al., 2012). Because muscle function is dependent on hemolymph [K^+^], muscle function can thus be restored before organismal osmotic homeostasis is fully re-established (Andersen and Overgaard, 2019; MacMillan et al., 2012).

The ionoregulatory model of chill susceptibility described above appears to apply to chill susceptible insects that are exposed to and rewarmed following a single, continuous cold period. In natural environments and in many conditions of insect storage for industrial applications, however, insects are subjected to frequent temperature fluctuations which can cause them to repeatedly cross critical threshold temperatures (Beardmore and Levine, 1963; Colinet et al., 2018; Javal et al., 2016; Long, 1970; Marshall and Sinclair, 2012). The extent to which insect thermal performance and/or life history is impacted by crossing threshold temperatures depends on the rates of temperature change, the frequency of these events, and the magnitude of the extremes experienced (Colinet et al., 2015). In outside environments, insects experience naturally occurring fluctuations in temperatures, while in a lab setting, temperature variations can be, for example, produced by creating a Fluctuating Thermal Regime (FTR). These FTRs are characterized by long cold periods interrupted by brief warm episodes or vice-versa (Colinet et al., 2016, 2015; Lalouette et al., 2011) and are unnatural in the sense that they don’t mimic real world scenarios of cold and warm temperature magnitude or duration. They also focus on physiological repair mechanisms during the short warm period, which contrasts from other studies that focus on repeated exposures to a stressful temperature (Marshall and Sinclair, 2012). Nonetheless, the main advantage of FTRs is that they provide a survival benefit in insects that can be used to prolong their “shelf life” for storage purposes (Chen and Denlinger, 1992; Colinet and Hance, 2010; Rinehart et al., 2011).

If cold survival is related to the ability to maintain and/or restore low hemolymph [K^+^] during and/or following a cold stress, FTR and repeated stress exposures may exert their fitness benefits through mitigating hyperkalemia and its downstream effects on cell survival. In keeping with this idea, adult fire bugs (*Pyrrhocoris apterus*) and beetles (*Alphitobius diaperinus*) in an FTR suffer less chilling injury and maintain lower hemolymph [K^+^] than individuals of the same species held at a constant low temperature (Koštál et al., 2007). Similarly, quiescent *D. melanogaster* larvae in an FTR exhibited reduced chilling injuries and maintained lower levels of extracellular [K^+^] compared to larvae exposed to a continuous low temperature (Koštál et al., 2016). Larvae in the same FTR also better maintained energy balance (Koštál et al., 2016). Together, these studies demonstrate that fluctuating temperatures are beneficial to survival compared to constant cold exposure, and this effect is at least partly attributable to better maintenance of ion homeostasis in insects. While some FTR studies involved a difference in the total amount of time spent in the cold between the constant and fluctuating temperature groups (Koštál et al., 2007; Petavy et al., 2001), more recent approaches attempt to match the cold duration of all treatment groups in a study that operates based on repeated stress exposures. Like in an FTR treatment, adult flies that experience repeated cold stresses also exhibit higher survival compared to individuals that persist in a sustained cold environment, but this survival benefit comes at the cost of lower fecundity (measured as the intrinsic rate of population increase) (Marshall and Sinclair, 2010). A similar trade-off was also observed in the goldenrod gallfly (*Eurosta solidaginis)*, where individuals that were subjected to repeated cold stresses yielded fewer offspring that had lower body masses (Marshall and Sinclair, 2018). Female *Drosophila suzukii* that were exposed to repeated cold stresses also produced fewer offspring (although offspring size was not measured) (Enriquez et al., 2018). A likely explanation for this apparent trade-off between survival and reproduction is that repeated crossing of threshold temperatures induces a re-allocation of energy stores (e.g. toward cryoprotection or restoration of metabolic or ionic homeostasis) that improves survival, but comes at the cost of reproduction during the long warm periods (Marshall and Sinclair, 2010).

In this study, we hypothesized that periods of warmth immediately initiate the process of homeostatic recovery after a cold exposure, while full recovery may take many minutes to hours. We thus predicted that even very short warm periods in between cold exposures could lead to increased survival and faster chill coma recovery relative to sustained cold stress. If these short warm periods were too rapid to permit complete restoration of ion and water balance, we expected that longer warm periods would permit greater survival and faster recovery than very brief warm periods. Given that recovery from chill coma has an apparent metabolic cost (MacMillan et al., 2012), and in keeping with prior evidence from *D. melanogaster* (as discussed above), we predicted that these improvements in survival as a result of warm periods would come at the cost of reproductive output from female flies in the days following the cold stress.

## Methods

### Animal husbandry and experimental design

The line of *Drosophila melanogaster* used in this study are derived from isofemale lines collected from a London, and Niagara on the Lake, Ontario (Marshall and Sinclair, 2010). Flies were reared at 22.0 ± 0.2°C on a 12:12 h (L:D) cycle and were fed a banana-based diet (primarily banana, corn syrup, agar, and yeast). In preparation for the experiment, groups of ∼200 adult flies were moved to a fresh bottle of food (180 mL bottles containing 50 mL of diet) for 2-3 h to lay eggs. Adult flies were collected on the final day of emergence and transferred to fresh food vials (40 mL vials containing 7 mL of diet), where they remained for three days. Female flies were collected under brief and light CO_2_ anesthesia, transferred to new food vials in groups of ten, and given two additional days to recover from the CO_2_ exposure before starting experiments to avoid known physiological effects of CO_2_ (MacMillan et al., 2017; Nilson et al., 2006). For most experiments, all female flies were sexed within 24 h of emergence, and all flies were five days old at the time of the experiment and were assumed to be mated. For the fecundity experiment, virgin flies were collected under brief CO_2_ anesthesia and collected within 12-14 h of emergence and kept in fresh food vials for five days to ensure they had not and could not mate.

On the day of the experiments, flies were blindly assigned to one of four experimental groups: (i) a Control (CTL) group which was held in the incubator at 22°C in fresh food vials for 25 h; (ii) a Long Cold (LC) group kept in the ice bath at 0°C for 25 h; (iii) a Repeated Cold 120 (RC120) group kept in the ice bath for 5 h then moved to the incubator for 120 min at 22°C, repeated five times in total; and (iv) and a Repeated Cold 15 (RC15) group kept in the ice bath for 5 h then moved to the incubator for 15 min at 22°C, repeated five times in total (Figure 1). Cold stresses were applied with the flies held in otherwise empty plastic vials containing 10 flies and with a foam stopper holding the flies in the bottom 20% of the vial. Flies were moved to vials containing food and placed in the incubator during the recovery periods. For the CCRT experiment, only LC, RC120, and RC15 were used, since control flies were not in coma and effectively have a CCRT of zero.

**Figure 1.**
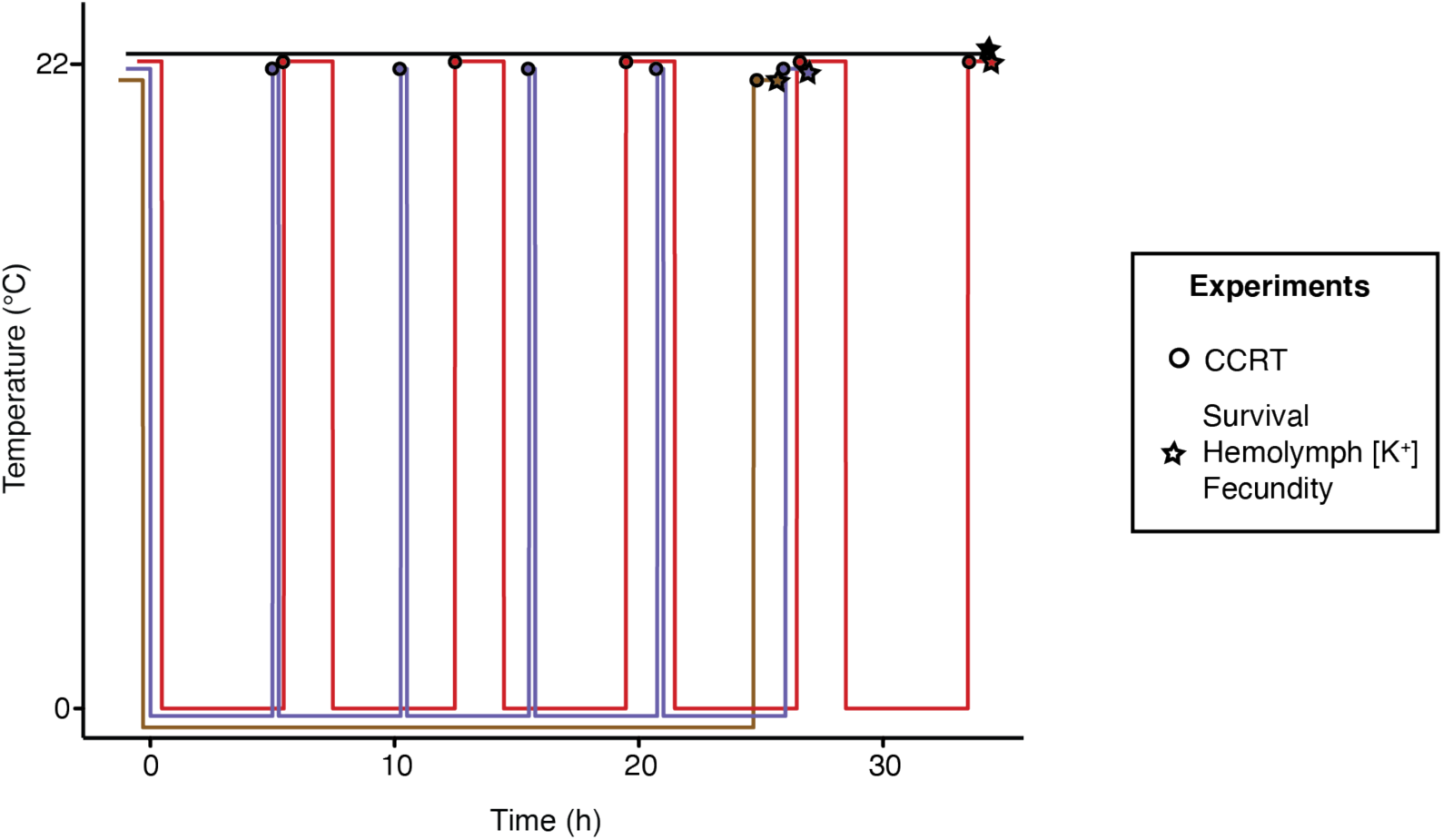
Duration and patterns of treatments applied to Repeated Cold 120 (RC120; red), Repeated Cold 15 (RC15; blue), Long Cold (LC; brown) and control (black) experimental groups. Both RC groups experienced 5 h at 0°C during each cold cycle, but RC15 spent only 15 min recovering at 22°C in between cold stresses while RC120 spent 120 min recovering. LC spent the full 25 h at 0°C. Lines slightly offset in both the x- and y-axes for clarity.

### Low temperature injury/survival assay

After a group had accumulated 25 h of exposure to 0°C, the flies were moved to food vials and placed in the incubator for 24 h. The degree of injury/survival of each fly was then assessed using a modified four-point scale similar to that described previously (MacMillan et al., 2018). Briefly, a fly was scored as a 1 if it was dead; a 2 if it was able to stand upright but not walk; a 3 if it was able to walk on the food surface but not climb the vial walls; and a 4 if it displayed baseline, pre-chilling behaviour by jumping, walking, and climbing inside the vial. This survival assessment was then repeated three more times (every 24 h) for a total of four days.

### Chill coma recovery time

Before the start of each cold exposure in the RC120 and RC15 groups, 18-20 individuals were taken out of the vials from each group and placed individually in small empty (4 mL) screw-thread vials. These vials were then placed in a small sealed bag and submerged in an ice slurry (roughly 1:3 ratio of water to ice) to create a 0°C environment in a Styrofoam box. Flies that were yet to undergo their respective cold cycle were kept in the standard *Drosophila* vials as in the previous experiment. At the end of a cold cycle, the bag was taken out of the ice bath and the vials containing the individual flies were quickly arranged on a laboratory bench at 22°C. A timer was started, and all vials were watched continuously. If a fly was able to right itself in 1.5 h, the time was recorded; otherwise, it was considered injured or dead. This procedure was done after each cold cycle in the RC120 and RC15 groups. For the LC group, 20 individuals were placed into the glass screw-top vials from the start and suspended in the ice bath for the full 25 h exposure.

### Fertility assay

We quantified offspring production following the methods of Marshall & Sinclair (2010). Immediately after the final treatment for a group of flies, 25 virgin females per treatment group were placed individually into new food vials with two untreated virgin males of the same age. After 24 h, each triad was transferred to a new food vial. This was done for a total of three times to yield four vials per triad collected over four days. The offspring were left to develop inside these vials, and total offspring production was measured by counting the number of adult individuals that emerged from pupae in each vial.

### Hemolymph [K^*+*^] measurements

After a group had finished its treatment, hemolymph was extracted from flies using the antennal ablation method previously described by MacMillan and Hughson (2014). During the hemolymph extraction from cold-exposed flies, vials with flies were held in an ice-water bath to keep the flies at 0°C. The extracted hemolymph was transferred into a dish containing silicone elastomer (Sylgard 184, Dow Corning Corporation, Midland, MI, USA). The dish was filled with hydrated heavy paraffin oil which fully covered the droplets, preventing evaporation. The concentration of K^+^ in the droplets was measured using ion-selective microelectrode technique (ISME) as described by MacMillan et al. (2015a). The ion-selective electrodes were made from glass capillaries (TW150-4, World Precision Inc., Sarasota, USA), pulled to a tip diameter of ∼3-5 µm using a P-1000 flaming/Brown micropipette puller. Once pulled the glass capillaries were heated to 300°C on a hot plate for 15 min before 20 µL of N, N-dimethyltrimethylsilylamine (Sigma Aldrich, Saint Louis, USA) was added using an inverted glass dish for 1 h. The capillaries were then cooled to room temperature, back-filled with a 100 mM KCl solution, and front-filled with a K^+^ ionophore cocktail (Potassium ionophore 1-cocktail B, Sigma Aldrich). Lastly, the ion-selective electrode tip was dipped in a solution of 1 mg mL^-1^ polyvinyl chloride (PVC, sigma: 81392) in tetrahydrofuran (Sigma Aldrich). The reference electrode was made by pulling a filamented glass capillary (USA 1B200F-4, World Precision Inc.) to a long and fine tip and backfilling it with 500 mM KCl solution.

Voltage was recorded using a ML 165 pH Amp and PowerLab 4/30 data acquisition system and Labchart® 6 Pro (AD Instruments, Colorado Springs, USA). Potassium concentration was determined by reference to two standards that differed in [K^+^] by a factor of ten (10 mM KCl, 100 mM KCl). The total ion concentration in the lower standard was controlled by replacing KCl with LiCl. Voltage values were converted to ion concentrations using equation (1):

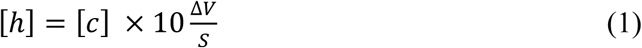

where [*h*] is the active concentration of K^+^ in the hemolymph, [*c*] is the concentration of K^+^ in one of the standard solutions, Δ*V* is the voltage difference between the standard solution and hemolymph, and *S* is the slope of the voltage response to the ten-fold concentration difference in the standard solutions. Voltage slopes were obtained from using a range of 50-60 mV from a ten-fold difference in ion concentration (MacMillan et al., 2015a).

### Data analysis

All data analysis was conducted in R v. 3.5.3 (R Core Team, 2019). Data distributions and variance structure were assessed using Shapiro-Wilk tests and Q-Q plots, and non-parametric approaches were used where appropriate and possible given the experimental design. Survival scores were analyzed among and within treatments using a linear mixed effects model using the lmer() function (in the lme4 package for R) with treatment and day (post cold stress) as factors and vial as a random effect (Bates et al., 2015). The relationship between initial mean survival score (on day 1) and the rate of decrease in survival score across the following three days was assessed using linear regression. A generalized linear model was used to test for an effect of treatment and the number of cold cycles (1-5) on chill coma recovery time. The effect of treatment on hemolymph [K^+^] was assessed using a Kruskal-Wallace test followed by pairwise Wilcox tests corrected for false discovery using the Benjamini-Hochberg (BH) method (Benjamini and Hochberg, 1995). The effects of treatment and egg laying date on cumulative offspring count was analyzed using a glm, and a Kruskal-Wallace tests followed by pairwise Wilcox tests (with BH correction) were used to test for differences in the total number of offspring produced on each egg laying date. Flies that died and/or produced no offspring at any point in the experiment were excluded from the analyses, so our fecundity data should be interpreted as the reproductive capacity of survivors, specifically. We saw no significant effects of treatment on the sex ratio of the offspring produced or the intrinsic rate of population increase (r; see supplementary data).

## Results

### Low temperature injury/survival

Periods of cold recovery strongly impacted survival outcomes in the four days following cold stress in *D. melanogaster*; there were significant main effects of cold treatment group (i.e. repeated cold exposures vs. constant cold exposure; LME; *F*_3,1483_ = 30.1, *P* < .0001) and the day of post-cold stress assessment (LME; *F*_1,1483_ = 10.1, *P* = 0.001; Figure 2A) on survival. Flies that received warm breaks (RC120 and RC15) had much higher rates of survival in the days following cold stress than those that experienced prolonged chilling (LC), and longer breaks were significantly better for survival that short breaks. Survival scores of chill injured flies also decreased in each subsequent day that they were assessed, with the LC group showing the largest decrease in survival scores over time (Figure 2A). This difference in the rate of change in survival scores drove a significant interaction between the effects of treatment and day of assessment on survival (LME; *F*_3,1483_ = 3.36, *P* = 0.018). This relationship was further examined using linear regression, which demonstrated that the mean survival score on day 1 (24 h following cold stress) was highly predictive of the rate of decline in survival scores over time (*P* = 0.020, R^2^=0.94; Figure 2B).

**Figure 2.**
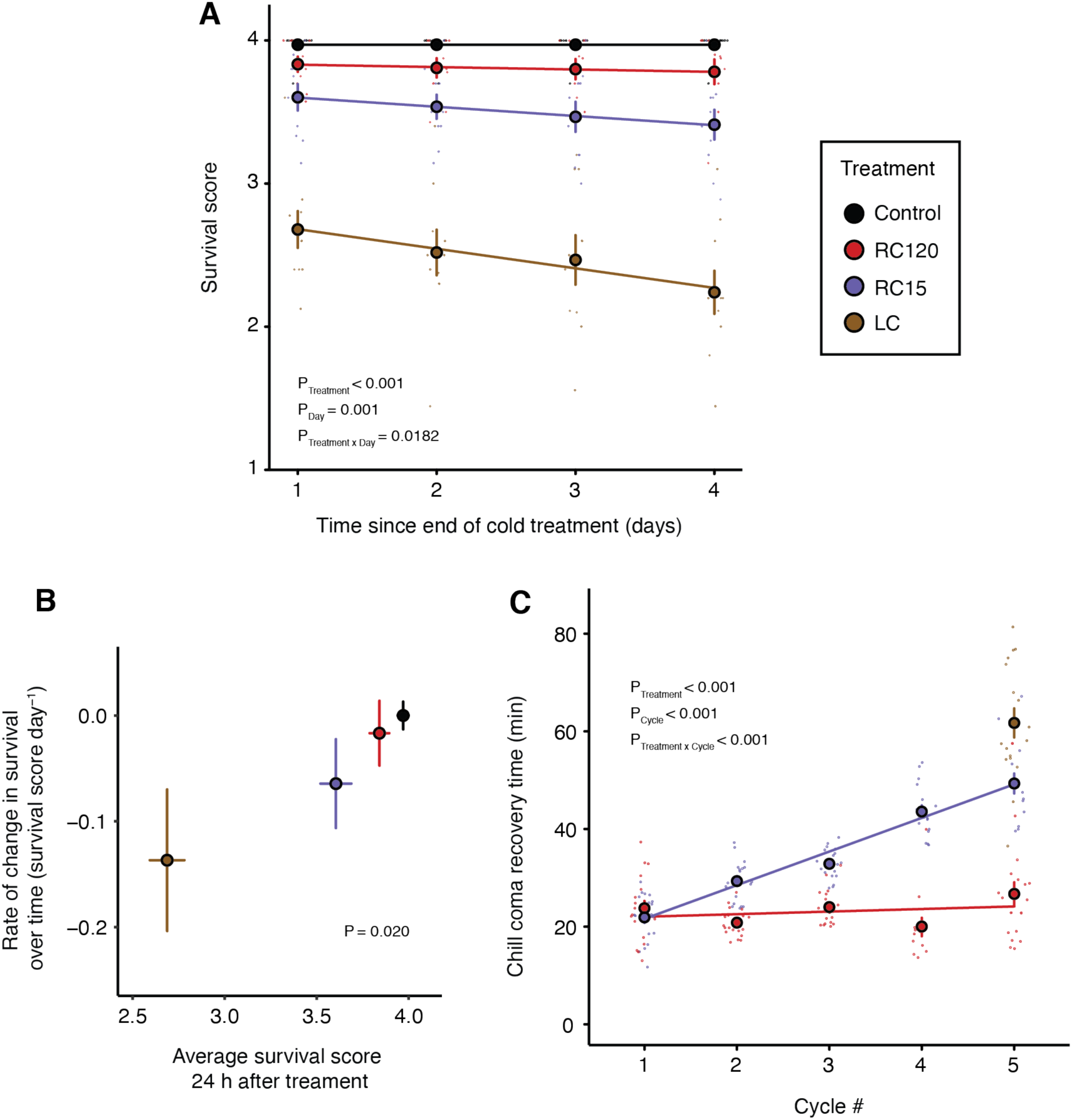
The immediate and latent effects of warm period duration on survival and chill coma recovery following repeated cold stress in female *D. melanogaster*. A) Survival scores of flies following constant (Long Cold, LC), repeated (Repeated Cold 120, RC120; Repeated Cold 15, RC15), or no (Control, CTL) cold stress. Flies were observed daily for four days to assess survival and scored on a scale from 1 to 4 (see Methods section for details of scoring method). Longer recovery times, as well as the day on which the survival assessment was made, significantly influenced survival of flies. n = 88-99 flies per experimental group. Small open circles represent individual data points and closes circles represent mean ± s.e. Error bars that are not visible are obscured by the symbols. B) Rate of decline in survival scores over the four days following the cold treatment, plotted against survival scores in thee first day of recovery. Continuous cold stress resulted in the greatest rate of decline in survival following the end of the treatment, followed by repeated cold exposures. C) Chill coma recovery time (CCRT) of flies in relation to the number of thermal cycles experienced. CCRT was measured in each cycle for both RC groups, and after the full 25 h at 0°C for the LC group. n = 13-20 flies for each cycle within each experimental group. Small open circles represent individual data points and closed circles represent mean ± s.e. Error bars that are not visible are obscured by the symbols.

### Chill coma recovery time

As with survival scores, the duration and frequency of cold recovery periods significantly affected CCRT in adult female *D. melanogaster*; recovery times were slowest in the LC group following 25 h of total cold exposure (61.7 ± 3.0 min), but were approximately 20% and 57% more rapid in the RC15 and RC120 groups, respectively (Figure 2C). In RC15, CCRT increased with each subsequent cold cycle, while there was no observed difference in CCRT between the first and fifth cold cycle in the RC120 group. This difference between the two groups experiencing repeated cold stress drove a strong significant interaction in the effects of cold cycle and treatment group on CCRT (GLM: *F*_1,184_ = 71.3, *P* < 0.001), meaning longer (2 h) recovery periods mitigated increases in CCRT across cycles, but CCRT increased progressively if recovery periods were shorter (15 min), and was maximized if no warm periods were provided (Figure 2C).

### Fertility assay

To test whether 15 or 120 min breaks in a cold stress have effects on female mating and/or offspring production, we used virgin females exposed the same treatment conditions as in the survival, CCRT, and ion balance experiments. Following their respective treatments, these females were given four consecutive days to lay eggs after mating. Periods of cold recovery (treatment group) strongly affected the total number of offspring produced (adults that eclosed) by female flies over four days (glm; *F*_3,244_ = 9.33, *P* < .0001), where the descending rank order of offspring production was control, RC120, RC15, LC (Figure 4A). There was also a significant interaction between the cold treatment and the rate of offspring production (ANOVA; *F*_3,244_ = 4.89, *P* = 0.003; Figure 4A). This interaction was specifically driven by the substantially slower rate of offspring production by the survivors of the LC treatment. Differences in the total number of offspring produced among the control and repeated cold stress (RC15 and RC120) groups appear to have been primarily driven by the RC flies tending to produce fewer offspring than the controls on the first day of egg laying (Figure 4B), but not on the following three days (Figure 4C-E).

**Figure 3.**
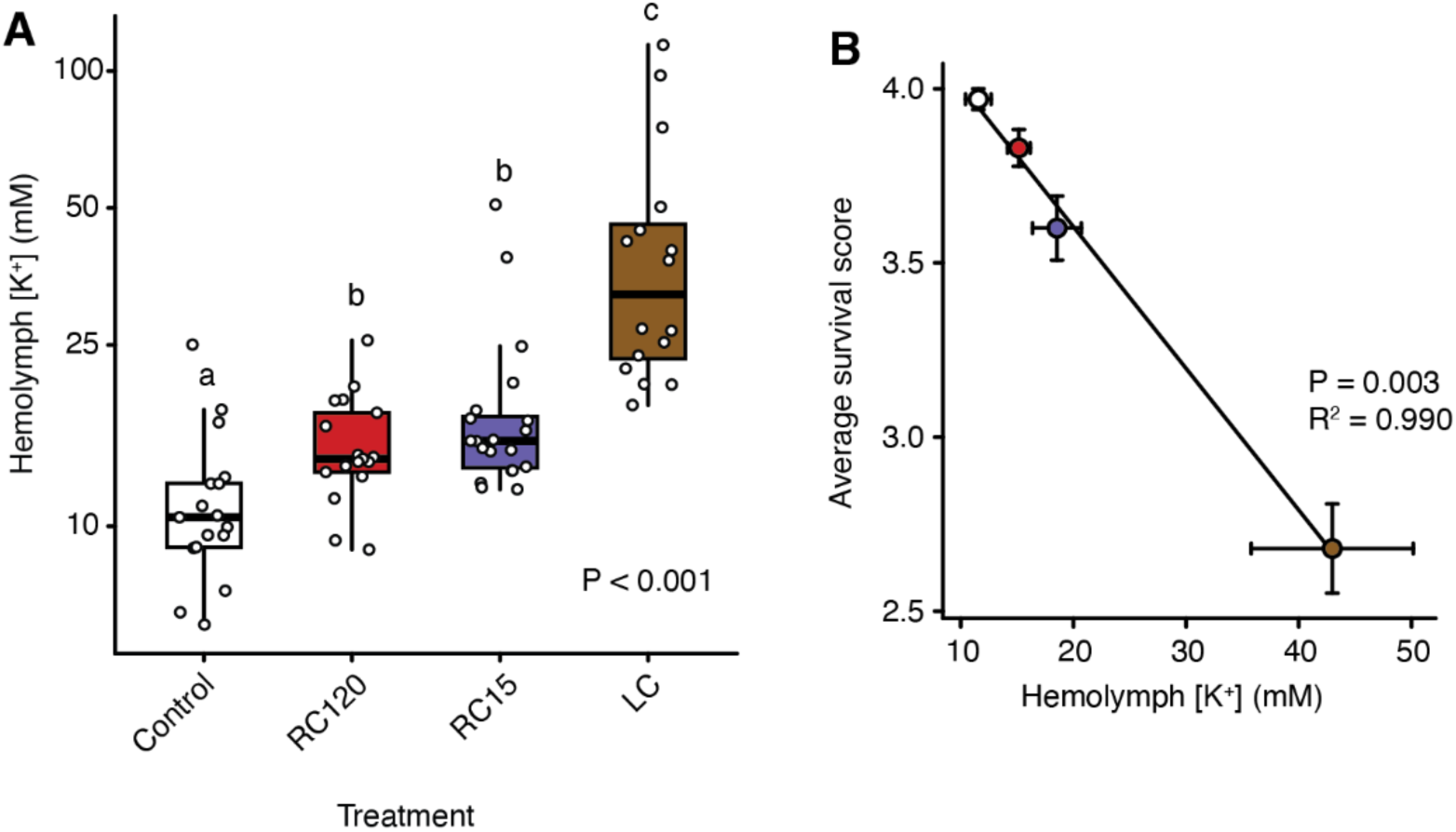
The effect of warm period duration on hemolymph [K^+^], and its relationship with survival following chilling, in female *D. melanogaster*. A) At the end of a group’s treatment, immediately upon being pulled out of the cold, hemolymph [K^+^] of flies that underwent constant cold stress (Long Cold, LC, brown), repeated cold stress (Repeated Cold 15, RC120, purple; Repeated Cold 120, RC120, red), or no cold stress (Control, CTL, white) was measured using ion-selective microelectrode technique (ISME). Both RC groups showed significantly lower [K^+^] compared to LC, but warm period length did not significantly affect [K^+^]. n = 16-20 flies per group. Small open circles represent individual data points. B) The average survival score of each group was plotted against their hemolymph [K^+^] to determine any relationship between the two. The regression analysis showed that the average hemolymph [K^+^] of each treatment group strongly predicted its average survival, shown by a strong negative correlation.

**Figure 4.**
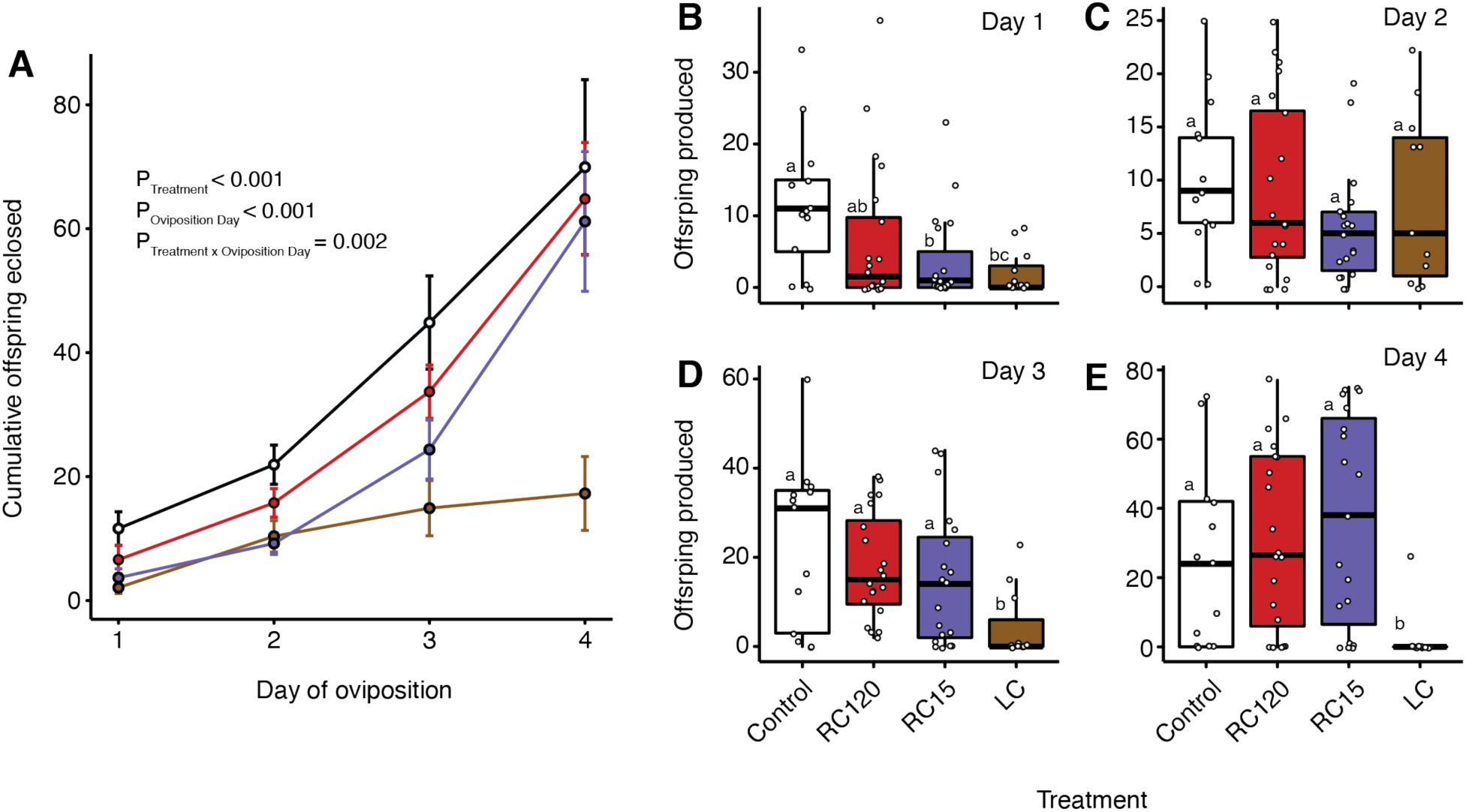
The effects of warm period duration on offspring production in female *D. melanogaster*. A) Following constant cold stress (Long Cold, LC, brown), repeated cold stress (Repeated Cold 15, RC120, purple; Repeated Cold 120, RC120, red), or no cold stress (Control, white), 25 virgin females from each group were placed in individual vials with two virgin males of the same age and left to mate for 24 h before being transferred to a new vial. This was done three times in total to yield four vials per triad. Offspring eclosions were counted among all vials until all eclosions ceased. Cumulatively, no significant difference in eclosion numbers were seen between CTL, RC120, and RC15. n = 11-20 flies per group. Circles shown are mean ± s.e. B-E) In addition to cumulative offspring counts, the amount of offspring produced on each day of egg laying were also counted to look for differences in day-to-day eclosion rates. Shorter warm periods led to lower offspring numbers only on day 1, compared to control flies. No differences in offspring counts between control, RC15, and RC120 were seen on the following three days. However, LC exhibited significantly lower offspring counts on day 3 and 4 due to parental death and lack of eggs. Open circles represent individual data points.

### Hemolymph K^*+*^ balance

To compare the effects of cold exposure treatments on the tendency of flies to suffer from hyperkalemia, we measured hemolymph [K^+^] and the end of each treatment (25 h of total exposure to 0°C). Cold exposure treatment strongly affected hemolymph [K^+^] (KW: χ^2^ = 38.9, *P* < 0.001); Flies that experience no warm periods (LC) reached the highest levels of hyperkalemia, control flies had the lowest hemolymph [K^+^], and flies that experienced interrupted cold spells (with 15 or 120 min breaks) had intermediate [K^+^] levels at the end of the treatment (Figure 3A). The average hemolymph [K^+^] strongly predicted average survival scores (data from the survival experiment above, as hemolymph sampling is destructive) of each group 24 h after the end of the cold stress (P=0.003, R^2^ = 0.990; Figure 3B).

## Discussion

Here, we show that even very short (15 min) warm breaks spaced within a prolonged (25 h) cold stress can yield considerable survival and reproductive benefits for a chill-susceptible insect. Our results showed increased survival in female *D. melanogaster* that experienced repeated cold stresses (with both 15 or 120 min breaks) compared to a continuous cold period, which closely agrees with evidence for the beneficial effects of warm periods found in other insect species (Colinet et al., 2006; Koštál et al., 2007; Petavy et al., 2001). These differences in survival between the RC and LC groups are consistent with the data observed by Marshall and Sinclair (2010) and Enriquez et al. (2018), which showed that repeated cold exposures result in increased survival compared to sustained cold exposures, and warm period length was positively correlated with survival. In our study, the average survival of each group was strongly predicted by the average hemolymph [K^+^] at the end of their respective treatments, suggesting that the differences in survival observed are at least partly attributable to differences in the severity of hyperkalemia.

The difference in survival between our RC120 and RC15 groups also correlated with the rate of change in survival over the four-day period post-treatment (while the flies were back in benign conditions). This means that RC15 (short breaks) individuals experienced a more severe decrease in survival over four days compared to RC120 (long breaks), and thus that longer breaks are still more beneficial than short breaks, even once the repeated stress has ended. This latent cold death, however, was most pronounced in the LC group where mortality increased from 6% (6/96 flies) (score of 1 on survival scale) on day one (24 h after treatment) to 28% (27/96 flies) by day four (96 h after treatment). Together, these findings suggest that the greater the injuries sustained during the cold stress (and the more severe the hyperkalemia), the more likely a fly is to suffer further injury or die in the following days.

Flies in the repeated cold groups showed greatly reduced CCRT compared to the sustained cold group. The duration of the recovery period very strongly determined the speed with which flies recovered from chill coma in later cycles; the RC15 group showed a progressive increase in CCRT up to the fifth cold cycle, while RC120 showed no significant increase in CCRT across cycles. The current model of chill coma recovery (Overgaard and MacMillan, 2017) states that a) CCRT increases with cold stress duration and b) CCRT after a cold stress depends on the rate at which ion homeostasis is re-established. Chill coma recovery in *Locusta migratoria, Gryllus pennsylvanicus*, and *G. veletis* has been linked with the re-establishment of K^+^ homeostasis, and in these species CCRT appears to be related to how fast ion homeostasis is re-established (Andersen and Overgaard, 2019; Des Marteaux and Sinclair, 2016; Findsen et al., 2014; MacMillan et al., 2014; MacMillan et al., 2012). In our study, the total cold exposure duration was the same across all three cold treatment groups. Our results thus suggest that 120 min warm periods provided sufficient time for ion and water homeostasis to be re-established before the next cold period occurred (MacMillan et al., 2012), while 15 min of recovery was insufficient. Although the RC15 group tended to have higher hemolymph [K^+^] than the RC120 group after a total of 25 h at 0°C, these two groups did not significantly differ. This suggests that, while 15 min warm periods led to a longer final CCRT compared to 120 minute warm periods, this difference was not exclusively driven by a greater capacity to maintain K^+^ homeostasis, and may be more related to an improved ability to restore low hemolymph [K^+^] during rewarming in the RC120 group. This suggestion is in agreement with the observation that rapid cold-hardening yields survival benefits by increasing the rate of recovery from hyperkalemia in locusts (Findsen et al., 2013)

Among the cold treatment groups, there were more offspring cumulatively produced by surviving flies in both the RC120 and RC15 groups, compared to the LC group; The LC group yielded the fewest offspring in the four days following the cold stress. This result was somewhat surprising, as it contrasts with prior work on repeated cold stress in *Drosophila*. In both *D. suzukii* and *D. melanogaster*, repeated cold exposures have previously resulted in significantly fewer offspring than were produced by an equivalent continuous cold group (Enriquez et al., 2018; Marshall and Sinclair, 2010). A likely reason for this difference is due to our different experimental design; both previous studies applied cold stresses on a near 24-hour cycle, unlike our design, and the flies thus had considerably more time for warm recovery. This additional time may provide flies with a more significant opportunity to alter their physiology. Such alterations might include gene expression and protein production (e.g. those involved in heat shock proteins) (Koštál and Tollarová-Borovanská, 2009), activation of catabolic pathways relevant to tissue repair or preparation for a subsequent stress (Wang et al., 2006), or initiation of reproductive diapause (Rinehart et al., 2007). A unified consequence of such responses would be the redirection of resources away from egg production, which would result in fewer offspring. Thus, while our fecundity results appear in contrast to those reported previously, we suspect that longer (e.g. ∼24 h) warm periods do yield a survival-fecundity trade-off, as suggested by Marshall and Sinclair (2010), while those experienced on shorter time scales (like ours) might not. In our design we did not see evidence for this trade-off, and we thus argue it is driven by adaptive (e.g. acclimation, quiescence) or maladaptive physiological responses to chilling that suppress fertility and which take considerable time, rather than physiological consequences of cold stress (e.g. injury) that occur while the stress is applied. From an applied perspective, this could mean that an “ideal” cycle may exist for a given species or population of insects that preserves fertility while also yielding survival benefits, but whether or not these patterns hold for a more extended period of insect storage remains unclear.

## Conclusion

In this study, even very short breaks in repeated cold exposures mitigated hyperkalemia and resulted in higher survival, lower CCRT, higher fertility in female *D. melanogaster* compared to sustained cold exposures. In addition, longer warm periods in between cold periods resulted in even higher survival and lower CCRT, but had no effect on offspring number, compared to shorter warm periods. These results indicate that differences in survival after repeated cold exposures of different warm period lengths do not necessarily arise as a result of differences in [K^+^] disruption but may be related to the time required to restore K^+^ balance during warm breaks. While this experimental design is unlikely to naturally-occur, fluctuating temperatures do occur over many different timescales in nature. While the short breaks applied in our experiments seem to result in both greater survival and fecundity in *D. melanogaster*, prior evidence suggests that this is not the case when treatments are applied over more ecologically-relevant timescales, and that trade-offs between these two traits may emerge only when there is sufficient time at benign temperatures for integrated physiological responses that divert resources away from reproduction.

## Supporting information

Data archive

## Acknowledgements

We wish to thank Bassam Helou, Justine Brown, Ravneet Hansi, Kaylen Brzezinski, and Dawson Livingston for their assistance with data collection throughout this study.

## Data Availability

All data is provided as a supplementary file for review and the same file will be uploaded to a data repository (e.g. Dryad) should the manuscript be accepted for publication.

